# Frustration and Fidelity in Influenza Genome Assembly

**DOI:** 10.1101/636613

**Authors:** Nida Farheen, Mukund Thattai

## Abstract

The genome of the influenza virus consists of eight distinct single-stranded RNA segments, each encoding proteins essential for the viral life cycle. When the virus infects a host cell these segments must be replicated and packaged into new budding virions. The viral genome is assembled with remarkably high fidelity: experiments reveal that most virions contain precisely one copy of each of the eight RNA segments. Cell-biological studies suggest that genome assembly is mediated by specific reversible and irreversible interactions between the RNA segments and their associated proteins. However, the precise inter-segment interaction network remains unresolved. Here we computationally predict that tree-like irreversible interaction networks guarantee high-fidelity genome assembly, while cyclic interaction networks lead to futile or frustrated off-pathway products. We test our prediction against multiple experimental datasets. We find that tree-like networks capture the nearest-neighbor statistics of RNA segments in packaged virions, as observed by EM tomography. Just eight tree-like networks (of a possible 262,144) optimally capture both the nearest-neighbor data as well as independently measured RNA-RNA contact propensities. These eight do not include the previously-proposed hub-and-spoke and linear networks. Rather, each predicted network combines hub-like and linear features, consistent with evolutionary models of interaction gain and loss.

---

The influenza virus is unusual in having a segmented genome, spread across eight RNA strands [1]. The negative-sense genomic RNA is transcribed in an infected cell’s nucleus to form positive-sense RNA, which undergoes both translation (to synthesize viral proteins) as well as replication (to form new genomic RNA). Segmentation allows genomic re-assortment, contributing to the emergence of novel influenza strains [2], [3]. However, segmentation also complicates the assembly and packaging of the complete viral genome into new virions [4]. Genomic RNA strands associate with specific viral proteins (nucleoprotein NP, and the polymerase complex PB2, PB1 and PA) to form rod-like viral ribonucleoprotein segments (vRNPs). Over 10,000 vRNPs are synthesized within four hours post-infection; these are packaged into nascent viral capsids at the plasma membrane, generating over 1,000 virions per hour [5]. Since each vRNP segment encodes essential proteins, all eight segments must be assembled and packaged to generate an infectious virion [1], [6]. Electron microscopy (EM) and fluorescence in-situ hybridization studies have shown that over 80% of new virions contain the complete genome, with each vRNP present in precisely one copy [7]–[9].

How does the influenza virus assemble its genome with such high fidelity? Genome assembly takes place as the vRNPs are trafficked to the plasma membrane [1], [4]. The selective packaging model [6], [7], [10], [11] posits that vRNPs bind to one another non-randomly via specific RNA-RNA and RNA-protein interactions; it is the resulting vRNP clusters that are packaged into virions. Consistent with this idea, mutations in the conserved RNA terminal regions cause defects in genome packaging [12]–[16]. Genomic RNA strands are seen to physically bind in vitro and in vivo [17]–[19] via RNA base-pairing interactions [20]–[22] and interactions mediated by vRNP-associated proteins [23]–[25]. In addition to the equilibrium RNA-RNA binding measured in vitro, live-cell imaging reveals that some inter-segment interactions are irreversible on the timescale of an infection [26]. Distinct vRNP segments are seen to co-localize inside the cytoplasm of infected cells [26], [27]. EM tomography of virions shows that the eight vRNPs are arranged as parallel rods, with electron-dense regions that suggest tight lateral interactions [19].

These studies strongly support the existence of specific interactions between vRNP segments. However they do not reveal the core interaction network that primarily drives genome assembly. Many possible interaction networks have been suggested, including a hub-and-spoke network (with a central “master segment”) and a linear network (looping to form a “daisy chain”) [10]. To our knowledge none of these hypotheses have been rigorously tested against the measured interaction data. Here we approach this problem by first exploring the influence of the inter-segment interaction network on the fidelity of genome assembly. We focus on the irreversible interactions, which create key decision points between correct and incorrect assembly pathways. Reversible and non-specific interactions [28] can play a role in stabilizing vRNP clusters already formed via irreversible interactions; we do not consider them here. By combining theoretical considerations with experimental datasets of virion structure and RNA-RNA interactions, we identify a handful of specific inter-segment interactions as the primary drivers of high-fidelity viral genome assembly.

## RESULTS

### Routes to high-fidelity genome assembly

We first explore the dynamics of the selective packaging model, in which genome assembly is driven by specific inter-segment interactions. The efficiency of a self-assembly reaction is typically measured by its yield: the fraction of total input material that is correctly assembled. A better measure for our purposes is fidelity: the fraction of output clusters that contain precisely one copy of each of the eight genomic RNA segments. Fidelity corresponds to the experimentally-measured fraction of new budding virions that are infections, assuming that clusters are uniformly packaged into viral capsids.

Irreversible interactions correspond to energetically favourable contacts between specific binding sites on the vRNP segments; these interactions could be orientationally rigid or flexible. We assume binding sites are organized such that two vRNP segments of the same type cannot bind to one another, and a given type of vRNP segment can bind to at most one copy of a given other type of vRNP segment. To assemble eight vRNP segments we require a minimum of seven interactions. Networks with precisely seven interactions are acyclic (tree-like), while those with more than seven interactions must include cycles (closed paths). There are 8^6^ = 262,144 tree-like networks (oeis.org/A000272) and over 250 million cyclic networks (oeis.org/A001187) that could potentially connect eight vRNP segments. Given an interaction network, we can model genome assembly as a stochastic chemical reaction (Methods). We start with a pool containing all vRNP types in equal amounts. We then allow the growth of clusters through pairwise aggregation, mediated by specific interactions between vRNP segments belonging to each cluster. We assume all allowed aggregation reactions occur by mass action with identical rate constants. Once no further aggregation events are possible, we calculate the final fidelity of the assembly reaction.

Cyclic interaction networks comprise the vast majority of possible networks. If the interactions in such networks are orientationally flexible, cycles will drive the futile synthesis of long polymers (Fig. 1A; X-Y-Z-X-Y…). Such futile reactions can be prevented by making the interactions orientationally rigid: the desired cluster with one copy of each segment is then stable because its binding sites are all either occupied or occluded. However this introduces a new problem: once all the monomeric vRNP segments are depleted, the assembly reaction gets stuck at frustrated oligomeric states (Fig. 1B; even though Y and Z can aggregate, X-Y and X-Z cannot since both copies of X compete to occupy the same position). This type of frustration has been observed in a broad class of self-assembly processes [29]. One way to prevent this is to use a fixed order of assembly, by tuning the aggregation rates (rapidly make X-Y, and then slowly make X-Y-Z). However vRNPs appear to aggregate in many possible orders (though some might be preferred [26], [27]). In this situation cyclic interaction networks will always show low fidelity, due to vRNPs being trapped in futile or frustrated off-pathway clusters. In contrast, tree-like networks of irreversible interactions will always show 100% fidelity (Fig. 1C,D; all aggregates reach state Y-X-Z), even when there is no fixed order, independent of the rates of aggregation, and whether interactions are flexible or rigid. This is surprising since tree-like interaction networks locally resemble cyclic interaction networks. A simple proof (Methods) shows that these results are completely general for tree-like and cyclic networks, regardless of the specific network topology. This strongly suggests that the core interaction network which drives genome assembly should be tree-like.

**Fig. 1.**
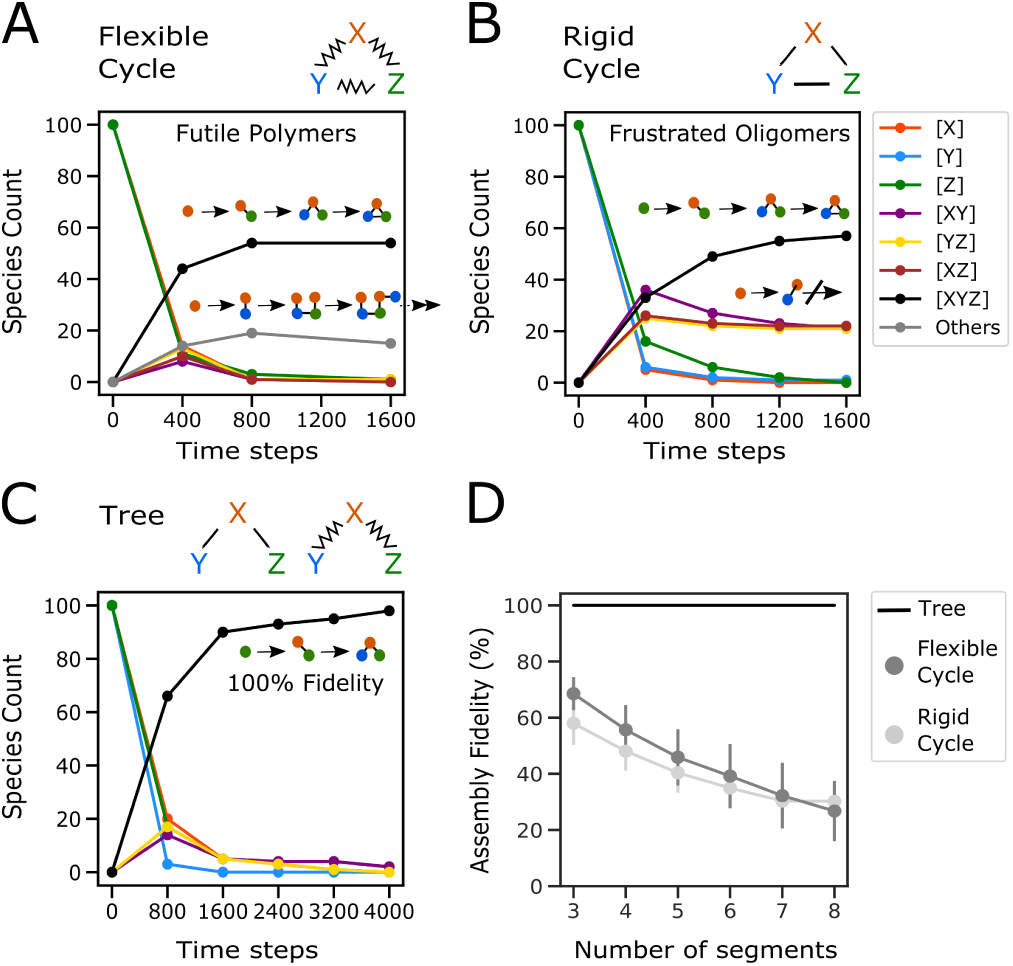
Stochastic simulation of genome assembly. We consider a toy model in which three segments (X, Y, Z) must assemble to form a desired cluster (XYZ). **(A**,**B**,**C)** Simulating time-dependent aggregation dynamics. The underlying interaction networks are indicated above each plot (zigzag edges show orientationally flexible interactions, straight edges show orientationally rigid interactions). The simulation starts with 100 copies of each segment and continues until no further reactions are possible. Counts of each possible reaction species over time from a single simulation are shown as curves of different colors (legend). Schematics show a few sample aggregation reactions, focusing on clusters that are present at long times. The fidelity is defined as the fraction of final clusters that are of the desired type (XYZ, black curves). **(A)** Flexible cyclic network. Flexible interactions allow the futile synthesis of long polymers. Segments are trapped within these futile clusters, reducing fidelity. **(B)** Rigid cyclic network. Once all monomeric segments are depleted, the remaining oligomers cannot bind to one another since identical segments compete to occupy the same spatial position. This is known as frustration. Segments are trapped within these frustrated oligomers, reducing fidelity. **(C)** Tree-like network. Once sufficient time has passed, all segments aggregate to produce the desired cluster XYZ, and no other cluster types are present. This reaction has 100% fidelity. **(D)** We consider systems with varying numbers of segments, whose interaction network is either a tree or a single long cycle. We compute the mean (± SD) fidelity over 500 stochastic simulations. Both flexible and rigid cycles show decreasing fidelity with increasing segment number. Trees always show 100% fidelity. These results generalise to all tree-like and cyclic networks, regardless of size and topology (proof in Methods).

### Inferring interaction networks from experimental data

EM tomography shows that the eight vRNP segments (henceforth numbered 1 to 8; Fig. 2A) are arranged in a characteristic “7+1” pattern within virions, with seven vRNPs on the periphery surrounding a central vRNP (Fig. 2B). Using vRNP length as a proxy for segment identity, all but the three longest segments (1, 2, 3) can be distinguished from one another [8]. The relative positions of the segments is found to vary from virion to virion, suggesting that interactions are orientationally flexible (Fig. 2B; see Methods for observed segment positions inside 30 virions [8]). However, certain vRNP pairs are more likely than others to appear as nearest neighbors. Segment identities can be further resolved by the SPLASH technique (Sequencing of Psoralen Crosslinked, Ligated, and Selected Hybrids) [30], which uses cross-linking and RNA sequencing to infer base-paired nucleotides in RNA complexes. SPLASH can be used to score the propensity of interaction between vRNP types in purified virions [20]. Both the EM tomography [8] and SPLASH [20] data are obtained for influenza strain A/WSN/33 (H1N1) in MDCK cells, allowing them to be directly compared.

**Fig. 2.**
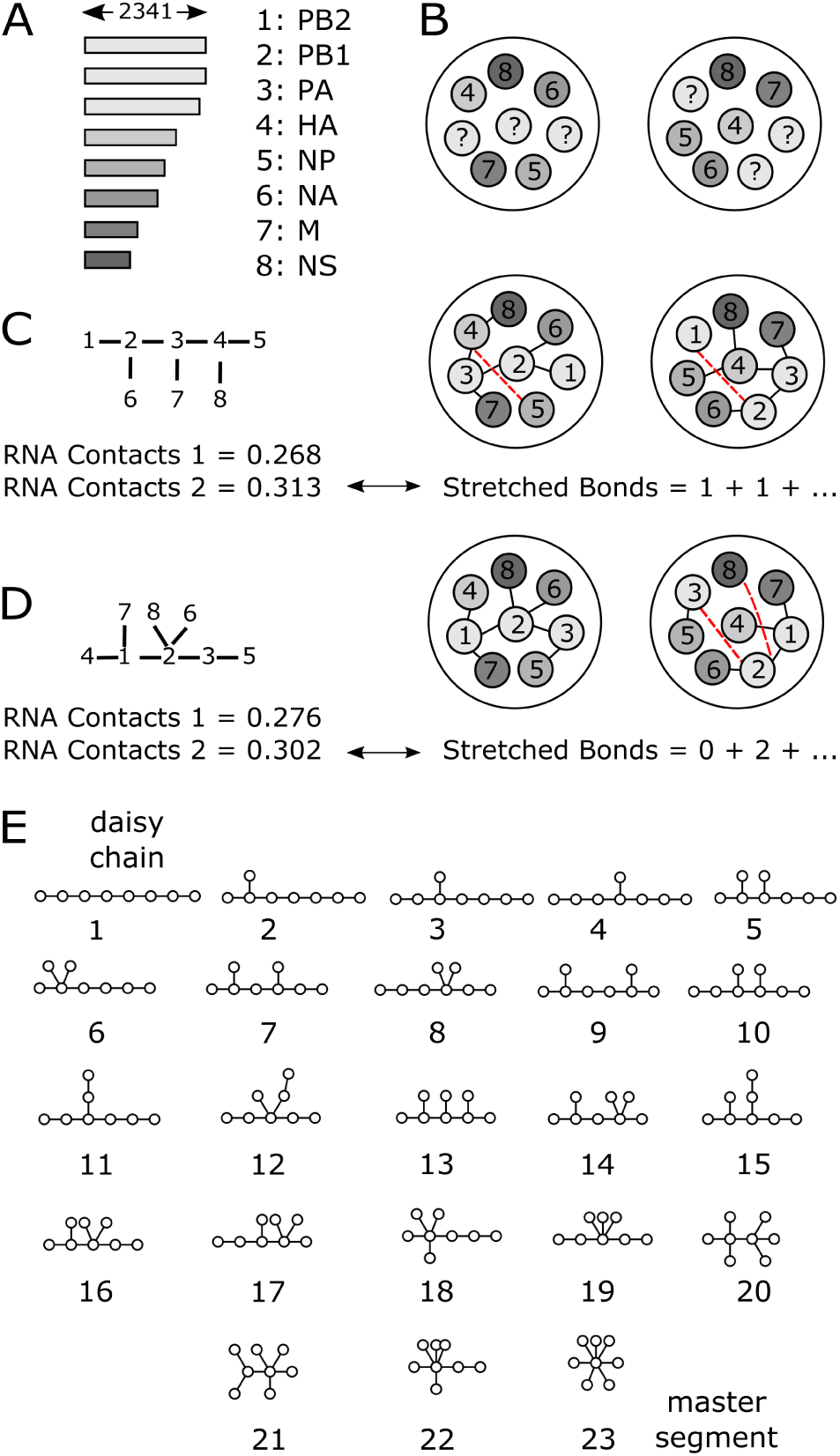
Genome packaging and inter-segment interaction networks. **(A)**. The influenza genome is made up of eight distinct RNA segments. Bar lengths show the number of nucleotides on each segment. Segments are conventionally numbered in order of decreasing length, and named according to the viral proteins they encode. **(B)** Each RNA segment is associated with viral proteins (NP, PB2, PB1 and PA) to form a rod-like viral ribonucleoprotein (vRNP) segment. Segments are packaged within virions as parallel rods in a “7+1” arrangement (as seen along the rod axes). EM tomography cannot distinguish between segments 1, 2 and 3 [8]; these are labeled “?”. The precise segment arrangement varies from virion to virion; two actual examples are shown here, out of 30 measured virions (Methods). **(C**,**D)** Given an interaction network (left) we assign it two types of scores. The ‘RNA Contacts’ score is the RNA-RNA interaction propensity measured by SPLASH [20], summed over each inter-segment bond in the network (two ‘RNA Contacts’ scores correspond to two SPLASH replicates). Higher ‘RNA Contacts’ scores indicate better agreement with the SPLASH data. The ‘Stretched Bonds’ score is the total number of stretched (non-nearest neighbor) bonds, summed across 30 virions observed by EM tomography. For each virion we use the assignment of segments 1, 2 and 3 that minimizes the number of stretched bonds (red lines). Lower ‘Stretched Bonds’ scores indicate better agreement with the nearest-neighbor data. **(E)** The 23 possible unlabeled tree-like network topologies for eight segments: Topology-1 is the linear network which forms a “daisy chain”; Topology-23 is the hub-and-spoke network with a central “master segment”.

Given an interaction network we assign it two types of scores (Methods). To obtain the ‘RNA Contacts’ score we simply sum SPLASH scores for all the bonds present in the interaction network (Fig. 2C,D: left; there are two scores corresponding to two SPLASH replicates). A higher ‘RNA Contacts’ score indicates better agreement with the SPLASH data. The ‘Stretched Bonds’ score is more involved, since there are six possible assignments of segments 1, 2 and 3 for each virion observed by EM tomography (Fig. 2C,D: right). For a given virion we select the assignment that permits the most bonds between nearest neighbors; to obtain the ‘Stretched Bonds’ score we then sum the number of stretched (non-nearest-neighbor) bonds across the 30 observed virions. A lower ‘Stretched Bonds’ score indicates better agreement with the virion nearest-neighbor data; an interaction network that captures all the observed nearest-neighbor occurrences would have a score of zero. Note that a cyclic network, compared to any tree-like sub-network, will have a better (greater or equal) ‘RNA Contacts’ score and a worse (greater or equal) ‘Stretched Bonds’ score.

We first calculated ‘Stretched Bonds’ scores for every possible cyclic and tree-like interaction network. The best overall network was a tree (Fig. 3A) with a ‘Stretched Bonds’ score of 34 (thirteen virions had two stretched bonds, eight virions had one, and nine virions had none). The 131 best networks were tree-like, while the best cyclic network had a ‘Stretched Bonds’ score of 44. We next generated 1,000 synthetic datasets by randomly shuffling the peripheral segments of each virion, and found the best overall tree-like network for each shuffled dataset (Fig. 3A). The single most common network was the hub-and-spoke network with a score of 48 (seen 737 out of 1,000 times); the lowest observed score was 42 (seen 3 out of 1,000 times), far worse than the best score of 34 for real virions. This proves that nearest-neighbor occurrences in real virions are highly non-random (*p* < 0.001) and consistent with a tree-like interaction network.

**Fig. 3.**
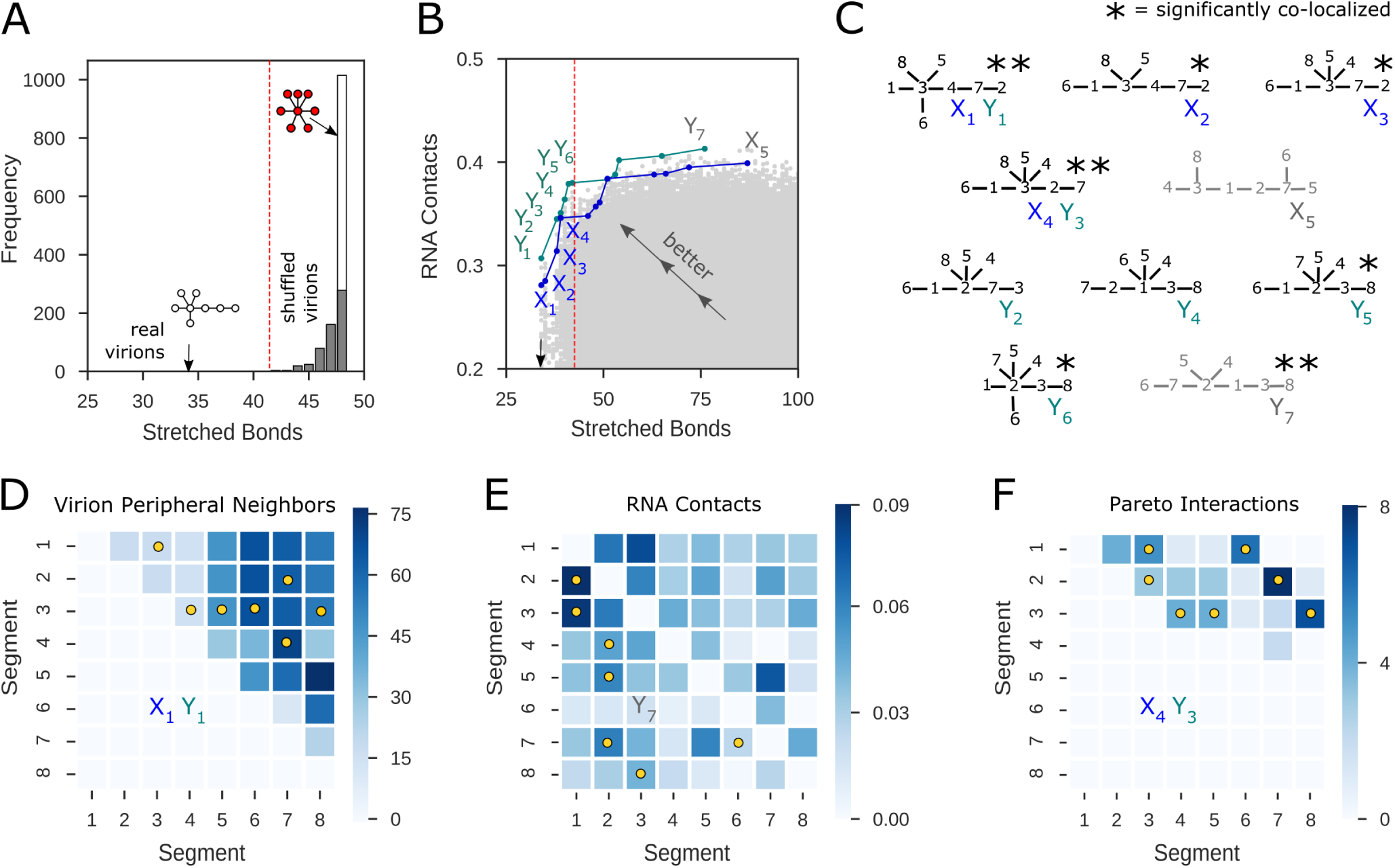
Inferring interaction networks from experimental data. **(A)** We calculated ‘Stretched Bonds’ scores (based on segment nearest-neighbor occurrences within 30 virions; Fig. 2C,D, right) for all possible interaction networks on eight segments (over 250 million networks). The best network was a tree (Topology-18; white network) with a score of 34. We repeated the same analysis for tree-like networks alone (262,144 networks) using synthetic datasets in which peripheral segments of each virion were randomly shuffled. The histogram shows the distribution of best scores obtained for 1,000 synthetic datasets, each containing thirty shuffled virions; the lowest score on shuffled data was 42 (seen 3 out of 1,000 times; the vertical red line shows this cutoff value). The white bar shows instances in which the best network was a hub-and-spoke (Topology-23; red network) with a score of 48 (seen 737 out of 1,000 times). **(B)** Scatter plot of ‘Stretched Bonds’ scores and ‘RNA Contacts’ scores for all 262,144 possible tree-like networks on eight segments. Each gray point corresponds to a single tree; each tree is represented by two points corresponding to two SPLASH replicates (Fig. 2C,D: left). Trees that are higher and to the left show better agreement with experimental data. The ‘Stretched Bonds’ score and ‘RNA Contacts’ score are jointly optimized by trees on the Pareto front, which form the upper-left envelope of the scatter plot. We show two Pareto fronts (blue, X labels; green, Y labels) corresponding to two SPLASH replicates. The black arrow shows the best ‘Stretched Bonds’ score of 34; the vertical red line shows the cutoff ‘Stretched Bonds’ score of 42. **(C)** The eight Pareto trees with ‘Stretched Bonds’ scores of 42 or less, corresponding to labels in Fig. 3B. We also show the two right-most trees on the Pareto front, which have the best ‘RNA Contacts’ scores (gray). Asterisks indicate trees whose bonds correspond to significantly co-localized segment pairs (two-tailed Mann Whitney U-test; * = significance value 0.05, Holm-Bonferroni method; ** = *p <* 0.005); co-localization *p*-values / 10^*-*3^, left to right [27]: 3.1, 7.0, 12.2, 1.3, 48.0, 195.0, 772.2, 5.9, 6.3, 1.3). **(D**,**E**,**F)** Inter-segment associations seen in different data sources. Yellow dots show inter-segment bonds in specific Pareto trees, indicated by their label from Fig. 3C. **(D)** Frequencies with which segment pairs are observed as peripheral nearest neighbors across 30 virions; each virion is represented by all six possible assignments of segments 1, 2 and 3, so the maximum possible score is 180. Since the data are symmetric, only the upper-triangular portion is shown. **(E)** Inter-segment RNA-RNA contact propensities measured by SPLASH. The upper-triangular and lower-triangular portions represent SPLASH scores for two different replicates. **(F)** Number of times each inter-segment interaction is observed among all eight Pareto trees with ‘Stretched Bonds’ score of 42 or less. Since the data are symmetric, only the upper-triangular portion is shown. The Pareto interaction map is much more sparse than the virion peripheral neighbor map and the RNA contact map.

For the remainder of our analysis we focus on the 262,144 possible tree-like networks. These fall into 23 topological classes (Fig. 2E): Topology-1 is the linear network (“daisy-chain”) and Topology-23 is the hub-and-spoke network (“master-segment”) [10]. We can represent each tree-like network as a point on a scatter plot (Fig. 3B), with the horizontal axis showing its ‘Stretched Bonds’ score and the vertical axis showing its ‘RNA Contacts’ score (points corresponding to the two SPLASH replicates are labeled X and Y). Trees that are higher and to the left dominate (i.e. are strictly better than) trees that are lower and to the right. Trees not dominated by any other tree, which jointly optimize the two scores, comprise the “Pareto front” (the upper-left envelope of the scatter plot). The leftmost tree on the Pareto front (X_1_/Y_1_, also shown in Fig. 3A) has the best-possible ‘Stretched Bonds’ score of 34 but a poor ‘RNA Contacts’ score. If we try to improve the ‘RNA Contacts’ score the ‘Stretched Bonds’ score gradually worsens until the shoulder value of 42 (Y_6_) past which it rapidly worsens. This (combined with the fact that 997 out of 1,000 shuffled datasets have ‘Stretched Bonds’ scores above 42; Fig. 3A) suggests we should only consider Pareto trees with ‘Stretched Bonds’ scores of 42 or less. There are only eight such tree-like networks (Fig. 3C), a massive reduction from the 262,144 initial possibilities. These eight Pareto trees have a median diameter of 4 interactions (compare with 2 for hub-and-spoke and 7 for linear) and a median max-degree of 5 interactions (compare with 7 for hub-and-spoke and 2 for linear). Segments 5, 6 and 8 are always tips; one among 1, 2 or 3 is always a hub. Across these Pareto trees, all but one interaction connect the set {1, 2, 3} with the set {4, 5, 6, 7, 8}.

### Evolution of interaction networks

Our theoretical model of high-fidelity assembly suggests the interaction network should be tree-like, but does not select a preferred tree topology. Genetic re-assortment studies show that inter-segment interactions evolve as viral strains diverge [2]. This process can make certain interaction network topologies more likely than others.

Consider first a simple model (Fig. 4A) in which a tree-like interaction network is grown by connecting new segments to randomly-chosen existing segments. The probability of obtaining a certain tree topology can be calculated by enumerating all possible growth orders starting from a single segment; this is equivalent to counting all consistent ways to label the segments of a given tree from oldest to the newest (Methods). Under this “Gain-only” model the probability is 4.0 × 10^*-*4^ for a hub-and-spoke network, 0.013 for a linear network, and 0.10 for the most probable Topology-8 network (Fig. 4C). A more realistic scenario is one in which interactions can be gained or lost (Fig. 4B). If we start with a viral population in which a given tree topology is fixed, a gain-plus-loss event can generate a new tree topology that has a chance of sweeping to fixation (we assume that all trees have equal fitness, all cyclic networks have low but non-zero fitness, and all disconnected networks have zero fitness). This can be modeled as a Markov chain whose equilibrium distribution gives the probability that the population has a given tree topology (Methods). Under this “Gain/Loss” model the probability is 2.2 × 10^*-*5^ for a hub-and-spoke network, 0.088 for a linear network, and 0.16 for the most probable Topology-3 network (Fig. 4C)

**Fig. 4.**
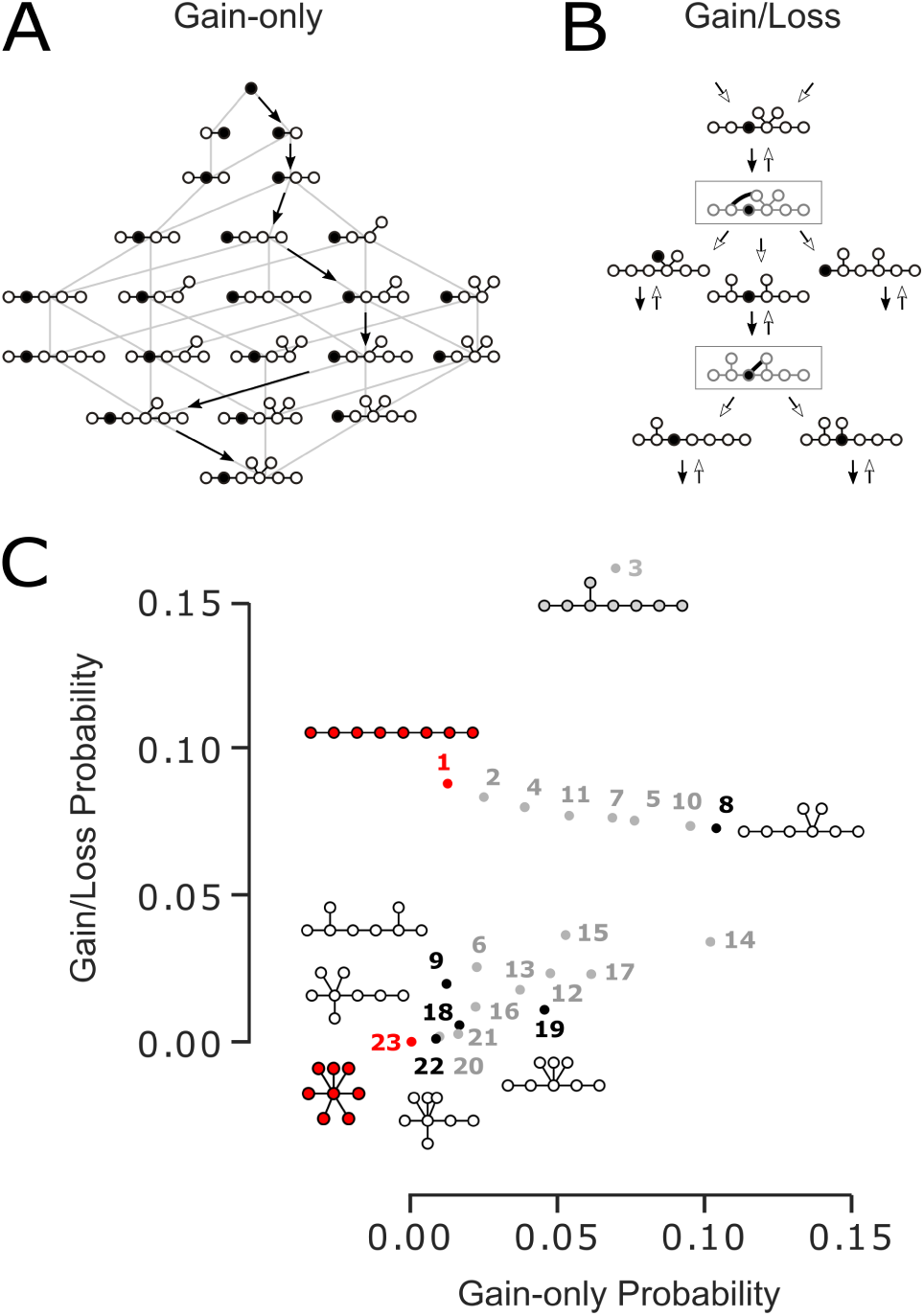
Evolution of tree-like interaction networks. **(A)** Under the “Gain-only model” new segments are attached to randomly-selected existing segments (black arrowhead = segment gain). The probability of obtaining given tree topology is calculated by summing all possible growth paths from all possible starting segments. In this example, there are 42 possible growth paths from the initial segment to the final tree. Similar calculations were be done for all distinct initial segments and final trees. **(B)** Under the “Gain/Loss” model trees evolve through repeated gain-plus-loss events (black arrowhead = interaction gain, white arrowhead = interaction loss), rapidly transition via cyclic states (gray boxes) to new tree topologies. This is a limiting case of evolution under successive population-genetic sweeps. It can be represented as a Markov chain that encodes transition rates between the 23 possible tree topologies, The probability of obtaining a given tree topology is found from the equilibrium distribution of the Markov chain. **(C)** Probabilities of tree topologies under the “Gain-only” and “Gain/Loss” evolutionary scenarios. Numbers correspond to the 23 tree topologies shown in Fig. 2E. We show the five topologies represented among the eight Pareto trees (white network and Fig. 3C). We also highlight the linear (“daisy chain”) and hub-and-spoke (“master segment”) topologies (red networks) as well as high-probability Topology-3 (gray network). The hub-and-spoke network is exceedingly unlikely under either evolutionary model.

While these abstract models cannot capture the evolutionary dynamics of real viral populations, they do make certain robust predictions. We can be confident that the highly-symmetric hub-and-spoke network is extremely unlikely to arise via the random gain and loss of interactions, unless it is specifically selected. More generally, we ought to expect interaction networks that combine both hub-like and linear features, rather either purely hub-and-spoke or purely linear networks. The five topologies represented among the eight Pareto trees are entirely consistent with this expectation (Fig. 4C).

## DISCUSSION

The mechanism by which the influenza virus packages its genome is a natural instance of a self-assembly process. There is growing interest in exploring the general principles of self-assembly across contexts: in complex multi-component biological systems such as ribosomes [31] and viruses [11]; and in synthetic systems such as colloidal aggregates and DNA tiles [32]. Kinetic assembly processes, in which aggregation reactions are driven out of equilibrium, allow greater control and higher yield of desired final products [31], [32]. However, out-of-equilibrium irreversible reactions can also lead the system down futile or frustrated paths [29]. Here we identify a simple design principle — a tree-like network of irreversible interactions — that provably achieves perfect fidelity.

Our approach is distinct from the challenge of finding the order of aggregation [27], which by definition is a tree whose nodes are clustered states and whose directed edges are reactions. The objects of our study are networks that could be cyclic or tree-like, whose nodes are segments and whose undirected edges are physical interactions. Our stochastic growth model (Fig. 1) does not select or require a single growth order, all growth orders consistent with the interaction network are permitted. The prediction that the network is tree-like then follows from the requirement of fidelity. This is due to a surprising property of tree-like interaction networks: they inexorably funnel growing aggregates toward the desired final product. Moreover this is achieved at the highest possible rate, since every aggregation reaction is productive (unlike equilibrium binding/unbinding reactions).

The purely theoretical preference for a tree-like interaction network allows us to extract useful information from the measured interaction propensities of vRNP segments. Since both our primary experimental datasets (EM tomography of nearest neighbors [8] and SPLASH RNA-RNA interaction measurements [20]) correspond to packaged virions, they cannot distinguish the irreversible interactions that drive genome assembly from weaker ones that stabilize the final assembled genome, or even incidental nearest-neighbor associations. This is why the virion peripheral neighbor map (Fig. 3D) and RNA contacts map (Fig. 3E) are both fairly dense. By requiring the interaction network to be tree-like and focusing only on Pareto trees, we remove such false positives to predict a sparse set of core interactions (Fig. 3F).

We must still ensure this procedure produces biologically-relevant predictions. We reasoned that segment pairs predicted to directly interact are more likely to co-localize during infection. The co-localization of all 28 segment pairs has been measured at eight hours post-infection [27]; this allows us to compare the co-localization of seven interacting segment pairs in a given tree with that of the remaining 21 segment pairs. We find that six of the eight Pareto trees are significantly co-localized (two-tailed Mann Whitney U-test, Holm-Bonferroni correction for multiple comparisons, significance level 0.05; Fig. 3C). This provides strong independent support for our prediction that the core interaction network is tree-like. However, co-localization depends on the time post-infection and potentially reports on direct as well as indirect interactions. Mutational studies could directly probe interacting regions, such as by swapping packaging signals between different RNA segments [15]. For example, the virus grows poorly when packaging signals of segments 1, 3, 5 and 7 are replaced with that of segment 6, but grows well when the same swap is done for segments 2, 4 and 8 [15]. This is most consistent with Pareto tree X_3_, in which segments 1, 3 and 7 are internal while the remaining segments are tips. In the predicted Pareto trees, almost all direct interactions involve segments 1, 2 or 3 (which encode vRNP-associated polymerase proteins PB2, PB1 and PA); whereas there are almost no direct interactions between segments 4, 6 and 7 (which encode capsid proteins HA, NA and M). This could enhance the re-assortment of the immunogenic capsid protein varieties between different influenza strains [3].

Further studies are needed to select the correct tree from among the predicted Pareto trees (and potentially others as well). We will need a principled approach to incorporate information from disparate data sources. Nevertheless, multiple lines of evidence (measurements of segment co-localization [27]; non-random nearest-neighbor propensities in packaged virions [8]; viral growth rates upon swapping packaging signals [15]) support our hypothesis that the core interaction network underlying influenza genome assembly is tree-like. These results not only provide insight into the dynamics of infection, but also have implications for understanding how new influenza strains emerge via genomic re-assortment and evolution.

## METHODS

### Stochastic simulation of genome assembly

We model genome assembly using a Monte Carlo simulation with discrete time steps. An interaction network on M segment types is specified, with either orientationally flexible or orientationally rigid bonds. We assume two segments of the same type cannot bind to one another, and a given segment type can bind to at most one copy of a given other segment type. We initialise the simulation with N copies of each segment type. As the simulation proceeds the segments irreversibly aggregate into clusters. Within each cluster we form satisfied bonds between every pair of segments that can interact. At each time step we select two of the clusters at random. We aggregate them into a single large cluster if a pair of segments, one from each cluster, has an unsatisfied bond. For the rigid bonds case we must also check that the two clusters do not both include the same segment type, since these would compete to occupy the same spatial position. We continue the simulation until no further aggregation events are possible. The final fidelity is then calculated as the fraction of clusters that are of the desired type, containing precisely one copy of each segment type.

### Proof that tree-like networks have 100% fidelity

The proof is by contradiction. We are given a flexible or rigid tree-like interaction network on M segment types, and given N copies of each segment type. The maximal cluster has exactly one copy of each of the M segment types. Suppose the final fidelity at the end of the aggregation process is less than 100%. There must be at least one cluster C with fewer than M segment types. Since the interaction network is connected, there must be at least one pair of segment types X and Y that interact, such that X is in C and Y is not in C. So the copy of X in C has an unsatisfied bond with Y. Since no further aggregation events are possible, every copy of Y either has a satisfied bond with a copy of X or (for the rigid bonds case) has an unsatisfied bond with X but belongs to a cluster C’ that cannot aggregate with C. This implies C and C’ both include some segment type Z. Any such Z is (directly or indirectly) connected to X in C and to Y in C’. Since we already know that X and Y can directly interact, this would mean the interaction network contains a cycle, which we know is not the case. Therefore no such cluster C’ exists. The only remaining possibility is that all N copies of Y have a satisfied bond with X. Since a given segment type can bind to at most one copy of a given other segment type, this implies that all N copies of X have a satisfied bond with Y, contradicting our assertion that there is at least one copy of X that has an unsatisfied bond with Y. This completes the proof.

### Proof that cyclic networks have less than 100% fidelity

We are given a flexible or rigid cyclic interaction network on M segment types, and given N copies of each segment type. To show that the average final fidelity is less than 100%, it is sufficient to show that there is at least one possible trajectory of the stochastic aggregation process with final fidelity less than 100%. First we treat the flexible case. Consider any length-L cycle in the interaction network, containing a pair of segment types X and Y that interact. We can grow two linear (L-1)-sized clusters C and C’ by single-segment-addition reactions so that C contains one copy each of all L segment types except Y, and C’ contains one copy each of all L segment types except X. C and C’ can then aggregate via the X-Y interaction to form a futile cluster containing two copies each of (L-1) segment types. This ensures the final fidelity is less than 100%. Next we treat the rigid case. Since a given segment type can bind to at most one copy of a given other segment type, the minimal length of a cycle is 3. Consider any length-3 cycle in the interaction network, containing segments X, Y and Z (the proof for a length-L cycle for any L ≥ 3 is identical). If N is odd, first form a single cluster XYZ by single-segment-addition reactions, leaving an even number of copies of each segment type X, Y and Z. If N is even, proceed to the next step. Let half of each segment type aggregate with its ‘left neighbor’ and the remaining half with its ‘right neighbor’ to form dimers XY, YZ, XZ. No further aggregation reactions are possible, since for any pair of dimers both will include the same segment type. The dimers are said to be frustrated. This ensures the final fidelity of at least one trajectory is less than 100%, so the average final fidelity is less than 100%.

### Modeling the evolution of tree-like interaction networks

Gain-only model (Fig. 4A): We are given segments labeled 1,…,N (this is an arbitrary label unrelated to vRNP segment identity). At each step of the process we take a tree with L segments and add segment L+1 to a randomly-chosen segment in the tree. We start with segment 1 and stop when we reach a tree with N segments. We record the final tree topology, ignoring segment labels. The statistical weight of a given tree topology under this process can be calculated as follows. List all distinct segment types, up to isomorphism (e.g. the hub-and-spoke topology has only two distinct segment types). Pick a segment type, root the tree at this segment and label it 1. Calculate the number of distinct ways, up to isomorphism, to label the remaining segments 2,…,N such that label values always increase along every branch. Summing this number over all distinct root segments gives the statistical weight of a given tree topology. To get the Gain-only probability we normalize this by the combined statistical weight of all possible tree topologies. Gain/Loss model (Fig. 4B): We model transitions between N-segment tree topologies as a discrete Markov chain. We are given a starting tree with N arbitrarily-labeled segments and (N-1) inter-segment interactions. There are (N-1)(N-2)/2 possible new inter-segment interactions. Adding a single new interaction gives a network with a single cycle. Removing any interaction in the cycle gives back a tree (e.g. removing the newly-added interaction gives back the original tree). Summing over all possible gain-plus-loss events and ignoring segment labels, we find the transition probability from the initial tree topology to any other tree topology. The Gain/Loss probability over tree topologies is the equilibrium distribution of this Markov chain.

### Scoring interaction networks

‘RNA Contacts’ score (Fig. 2C,D: left): From SPLASH measurements we obtain the total number of interactions observed between each segment pair [20]. We normalize this across all segment segment pairs to obtain an RNA-RNA contact propensity. The ‘RNA Contacts’ score of a given interaction network is the sum of contact propensities of the interacting segment pairs. ‘Stretched Bonds’ score (Fig. 2C,D: right): From EM tomography we obtain the arrangement of segments within packaged virions [8]. Segments 1, 2 and 3 cannot be distinguished by this method; for a given interaction network we find the assignment of 1, 2 and 3 within each virion that has the most interactions between nearest neighbors. The ‘Stretched Bonds’ score of the network is the sum of the number of stretched (non-nearest-neighbor) bonds across virions. The observed segment arrangements for 30 virions [8] are shown below. Each string represents a single virion, starting with the central segment and moving clockwise over peripheral segments; ‘?’ represents segments 1, 2 or 3.

4:87??65?, 4:86?57??, 4:857?6??, 4:85?6?7?, 4:8?75?6?,

4:8?75??6, 4:8?7?5?6, 4:8?6?75?, 4:8?6?7?5, 4:8?5?7?6,

4:8??765?, 4:8??75?6, ?:8754?6?, ?:8746?5?, ?:86547??,

?:8647??5, ?:86?57?4, ?:86?47?5, ?:8564?7?, ?:856?7?4,

?:856?47?, ?:8547?6?, ?:85?6?74, ?:847?6?5, ?:84657??,

?:8?674?5, ?:8?6?745, ?:8?546?7, ?:8?5?746, ?:8?4756?

## ACKNOWLEDGEMENTS

We thank M. S. Madhusudhan for critical feedback, and participants of the ‘Entropy, Information and Order in Soft Matter’ Program (ICTS/eiosm2018/08) at the International Centre for Theoretical Sciences for useful discussions.

## AUTHOR CONTRIBUTIONS

MT conceived the project. NF carried out the analysis. NF and MT wrote the paper.

